# Facultative parthenogenesis: a transient state in transitions between sex and obligate asexuality in stick insects?

**DOI:** 10.1101/2022.03.25.485836

**Authors:** Chloé Larose, Guillaume Lavanchy, Susana Freitas, Darren J. Parker, Tanja Schwander

## Abstract

Transitions from obligate sex to obligate parthenogenesis have occurred repeatedly across the tree of life. Whether these transitions occur abruptly or via a transient phase of facultative parthenogenesis is rarely known. We discovered and characterised facultatively parthenogenetic populations of the North American stick insect *Timema douglasi*, a species in which only obligately parthenogenetic populations were known so far. These populations comprised three genetic lineages. Females from all lineages were capable of parthenogenesis (with variable efficiency) but their propensity to reproduce sexually after mating varied extensively. In all three lineages, parthenogenesis resulted in the complete loss of heterozygosity in a single generation. Obligately parthenogenetic *Timema* have also lost all heterozygosity, suggesting that the transition to obligate parthenogenesis did not require a modification of the proximate mechanism, but rather involved a gradual increase in frequency. We speculate that facultative parthenogenesis may often be transient and be replaced by obligate strategies (either sex or parthenogenesis) because of a trade-off between the efficiency of the two reproductive modes. Such a trade-off could help explain why facultative parthenogenesis is rare among animals, despite its potential to combine the known benefits of sex and parthenogenesis.

## Introduction

The way organisms reproduce varies extensively across the tree of life. While most animals engage in some form of sex to produce offspring (Bell, 1982; The Tree of Sex Consortium, 2014), many species take alternative routes to reproduction. The most widespread alternative is parthenogenesis, whereby females produce offspring without genetic contribution from males (Normark, 2003; Suomalainen et al., 1987).

Most species appear to be either obligately sexual or obligately parthenogenetic in natural populations, even when there is some capacity for the alternative reproductive mode. For example, virgin females in many sexual species can on occasion reproduce through spontaneous parthenogenesis (“tychoparthenogenesis”), typically with very limited success (Bell, 1982; Fields et al., 2015; Liegeois et al., 2021). On the other end of the spectrum, recent studies have revealed rare or cryptic sex in several species previously thought to be obligately parthenogenetic (Boyer et al., 2021; Freitas et al., 2023; Kuhn et al., 2021; Laine et al., 2022). However, even though such species are still capable of sex, this mode of reproduction is used very rarely. Thus, species cluster at the extremes of the continuum and are either largely sexual or largely asexual (Bell, 1982). This is surprising, given that facultative parthenogenesis, whereby the capacity for sex or parthenogenesis of females are approximately equal, has the potential to combine the long-term advantages of recombination and segregation that come along with sex, together with the short-term advantages of asexuality (D’Souza & Michiels, 2010; Otto, 2009).

The relative rarity of facultative parthenogenesis also raises the question of whether obligate parthenogenesis generally evolves abruptly from sexual ancestors. This is the case for example in many or most obligately asexual lineages of hybrid origin (Cuellar, 1974; Schultz, 1973). Alternatively, the evolution of obligate parthenogenesis without hybridization may involve facultative parthenogenesis as an intermediate step, which would then be transient. We here evaluate these alternatives using the stick insect genus *Timema*.

*Timema* stick insects are native to the western USA and Mexico and have been studied extensively for their repeated transitions from sex to obligate parthenogenesis. Five of the 21 described species reproduce via female-producing parthenogenesis, while the others are sexual (Sandoval et al., 1998; Vickery & Sandoval, 1999, 2001). In parthenogenetic species, fertilization of oocytes does not occur even when females are mated with males of related sexual species in the laboratory (Schwander et al., 2013). However, cryptic gene flow has been documented in some species, presumably mediated by rare males (Freitas et al. 2023). Conversely, females of the sexual species are largely incapable of parthenogenesis, although spontaneous parthenogenesis in 1-2% unfertilized eggs laid by virgin sexual females is well documented (Schwander et al., 2010; Schwander & Crespi, 2009). When mated, sexual females fertilize all their eggs, and population sex ratios are close to 50:50 (Arbuthnott et al., 2015; Schwander et al., 2010). Thus the *Timema* populations described so far fit well with the general pattern in animals whereby reproductive strategies are largely obligate.

However, we recently discovered populations with a large proportion of males in the geographic range of the species *T. douglasi*, a species that was believed to be exclusively parthenogenetic. The male-containing populations were geographically close to female-only populations, with an apparently abrupt transition from female-only populations to populations with relatively even sex ratios. Such variable population sex ratios could, for example, result from mixes between different proportions of sexual and parthenogenetic females, or from facultative parthenogenesis with sex and parthenogenesis used at different frequencies. Alternatively, if the newly discovered populations were sexual, variable sex ratios could also stem from sex ratio distortion. For example, a driving X chromosome, which would be present in more than 50% of transmitted sperm cells, could also generate female-biased sex ratios (Helleu et al., 2015; Jaenike, 2001).

In order to investigate these different hypotheses, we sampled populations along two separate transects and quantified the local sex ratios. We then characterized the reproductive modes of females from populations with different sex ratios by measuring the capacity for parthenogenesis of virgin females and by distinguishing parthenogenetic and sexual offspring in controlled crosses via genotyping based on Restriction-Associated DNA Sequencing (RADseq). We also screened the sex ratio of sexual offspring for deviations from the expected 50:50 ratio to test whether a distorter could cause the observed biases in sex ratio. Finally, we also used the RADseq genotypes to assess whether females with different reproductive modes belonged to different genetic lineages, and to study the phylogenetic relatedness between these lineages and previously described lineages of parthenogenetic *T. douglasi* and close sexual relative *T. poppense*.

## Material and Methods

### Population sampling and reproductive mode characterisation

We characterized the population sex ratios at 29 sampling locations along two transects located in northern California, respectively referred to as “Manchester” and “Orr” transect (Fig. 1). For both transects, we chose the sampling locations according to the host plant distribution along the main road, with isolated patches of redwood (*Sequoia sempervirens*) or douglas fir (*Pseudotsuga menziesii*) considered as distinct locations. We sampled each location during 2 hours with approximately constant sampling intensity and collected up to 270 individuals (all juvenile) per location. We sexed the individuals based on external morphology. In total, we collected 1195 individuals for the Manchester transect (864 females and 331 males in 12 different populations, hereafter called Manchester 1 [westernmost] to Manchester 12 [easternmost]; see Figure 1) and 1074 individuals for the Orr transect (1029 females and 45 males in 12 populations, hereafter called Orr 1 [easternmost] to Orr 17 [westernmost]; no *Timema* were found at 5 locations).

**Figure 1.**
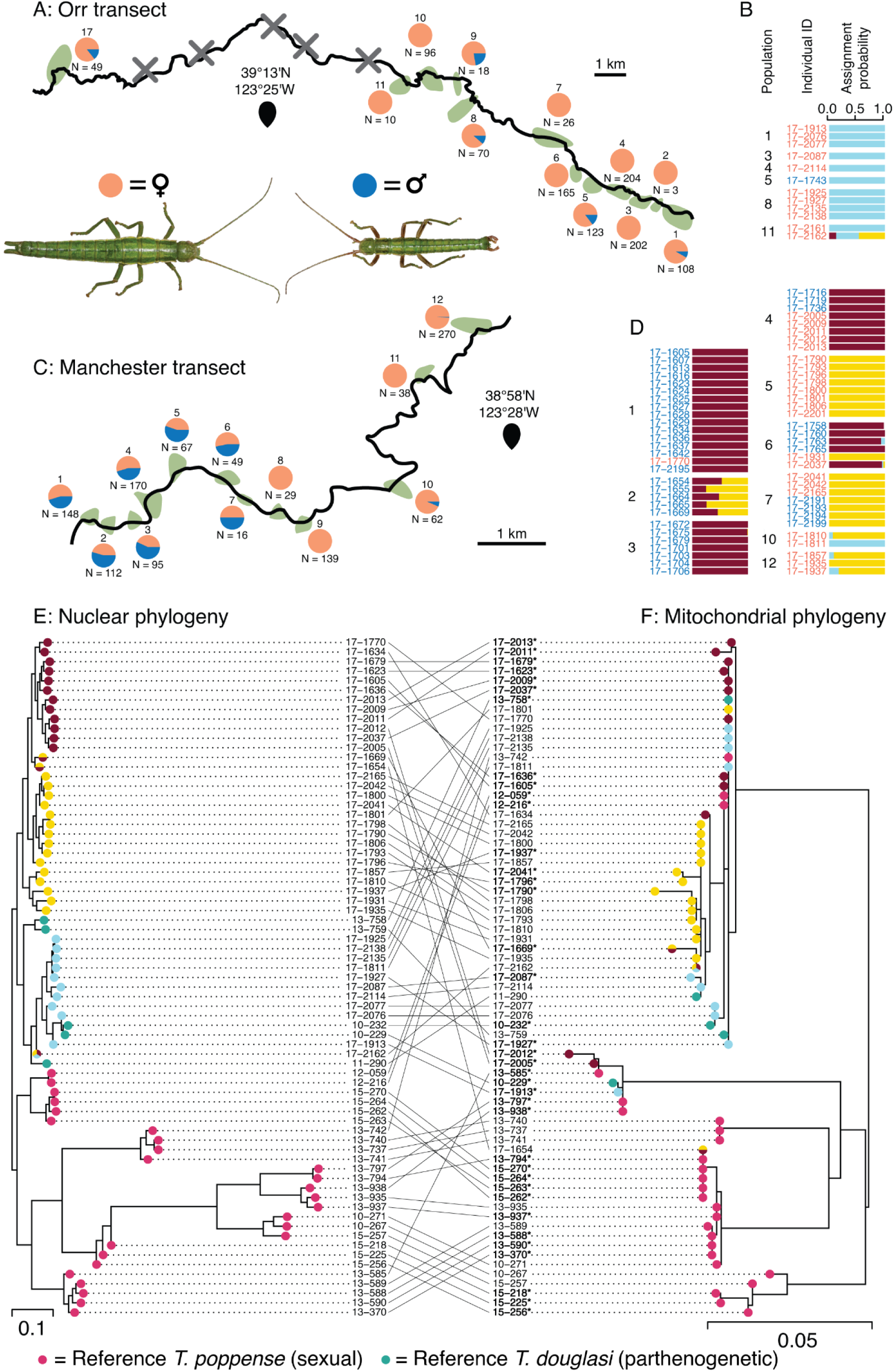
Sex ratios in the sampled populations, genetic structure and species identification of the collected individuals. **A** and **C**: Schematic maps of the Orr and Manchester transects, with host plant patches represented in green along the main road. Pie charts represent the sex ratio of individuals collected in the field. Crosses indicate surveyed host plant patches with no *Timema* present (locations 12-16 along the Orr transect). **B** and **D**: Assignment probabilities to each of the 3 genetic lineages for each genotyped individual, split by transect. **E**: ML nuclear phylogeny (16391 SNPs) of the genotyped field-collected individuals plus 26 *T. poppense* (sexual) and five *T. douglasi* (parthenogenetic) reference individuals to infer species assignments for individuals from the transects. **F**: ML mitochondrial phylogeny of the same individuals based on 159 polymorphic sites of the COI mitochondrial gene. One individual of *T. cristinae* was used to root the tree (not displayed). Tip colour refers to genetic lineage (individuals from Orr and Manchester transects) or reference individuals used for species assignment. Individuals denoted in bold with a star had double peaks in their COI sequence and the allele corresponding to the highest peak was retained. Timema photos taken by © Bart Zijlstra - www.bartzijlstra.com

We aimed to characterise the reproductive mode of approximately 10 females from each female-only population and of approximately 20 females from each population with both sexes (when available). However, the final numbers varied considerably because of mortality, and because females sometimes laid too few eggs prior to mating to determine hatching success (see below). Individuals of each population were separated by sex and maintained in cubic mesh cages (30 x 30 x 30 cm) on douglas fir, in a climatic chamber at 23°C, 55% humidity and 12:12h day:night cycle. Since all individuals were collected as juveniles and only adults mate, this allowed us to obtain virgin females for all populations. We isolated 352 females in Petri dishes (14.5 cm diameter) to obtain unfertilized eggs, with fresh food (a small branch of douglas fir), soil (for females to ingest in order to coat their eggs), and a moistened piece of cotton wool (to prevent desiccation) added every two days. Once a female had laid approximately 20 unfertilized eggs, we allowed her to mate with one to three males from sampling locations with high male frequencies (Manchester 1 - 4, 6 - 7, Orr 5). For each female, the eggs laid before and after mating were kept separately until hatching, which occurs after approximately 5-6 months of diapause. The eggs began to hatch on October 27th of the same year. The eggs were then checked every other day and the number of hatchings recorded. Hatchlings were stored in 100% EtOH until further use. No eggs hatched after December 27th.

In addition, we re-analysed previously published data on the capacity for parthenogenesis of seven sexual species (*T. petita, T. podura, T. cristinae, T. chumash, T. knulli, T. californicum* and *T. poppense*) from (Schwander et al., 2010; Schwander & Crespi, 2009) and characterized it for the five asexual species (*T. douglasi, T. shepardi, T. monikense, T. genevievae* and *T. tahoe*). These data represent a total of 450 virgin females (range: 3 – 162 per species) collected in 2007 and 2008. We reared them under the conditions described above, except that they were maintained at room temperature and fed with different host plants according to the natural host plant use of each species (see Schwander et al., 2010 for details). We collected all eggs and assessed the hatching success of all females which had laid at least five eggs (424 females, 14021 eggs).

### Genotyping

To test whether the individuals collected along the Manchester and Orr transects belonged to different genetic lineages, we genotyped 32 females and their mates (1 - 3 per female, 42 males in total). We also assessed whether eggs produced after mating were fertilized or not, and quantified heterozygosity in parthenogenetically produced eggs. To do this, we genotyped 3 hatchlings from eggs laid before mating (when available) for each of the 32 genotyped females (77 hatchlings from 27 females in total; for the 5 remaining females, none of the eggs laid prior to mating hatched), and 7 hatchlings from eggs laid after mating (when available) for 24 of the 32 females. For the eight remaining families, we genotyped all available hatchlings from eggs laid after mating (17 - 51), initially for a different project. In total, we genotyped 365 post-mating hatchlings. Details on the number of hatchlings and males per family are provided in Table S1.

We extracted the DNA of a leg (adults) or the whole body (hatchlings). We flash-froze the tissue in liquid nitrogen and ground it via sonication with ceramic beads. We then performed the extraction using the Qiagen DNA Blood and Tissue kit on a BioSprint 96 workstation, following manufacturer protocols.

We built double-digested RADseq libraries following the protocol by Brelsford et al. (2016). In short, we digested genomic DNA with EcoRI and MseI and ligated Illumina TruSeq adapters to the cut site. The EcoRI adapter included an individual 8-base barcode. We amplified the ligated fragments in 20 PCR cycles. We then multiplexed the libraries and size-selected fragments of 300 - 500 bp on an Agarose gel. The resulting libraries were then sequenced on 10 Illumina HiSeq 2500 lanes (100 - 125 bp, single end) at the Lausanne Genomics Technology Facility.

We built loci using Stacks version 2.3e (Catchen et al., 2013). We first demultiplexed the raw reads with the process_radtags module with parameters -c -q -r and --filter-illumina and trimmed all the reads to 92 bases. We then mapped the reads to the reference genome of *T. douglasi* (Bioproject accession number: PRJEB31411; Jaron et al., 2022) using the mem algorithm of BWA version 0.7.17 (Li & Durbin, 2010). We built loci using the gstacks module, enabling the --phasing-dont-prune-hets option. Finally, we output a vcf file with a single SNP per locus using the --write-single-snp option in populations. We then filtered genotypes in Vcftools version 0.1.15 (Danecek et al., 2011) as follows: we retained genotype calls if they had a minimum coverage depth of 11, and discarded loci that had a minor allele count lower than 3 or a genotype call in fewer than 75% of all individuals. We further discarded five individuals which had more than 60% missing genotypes. Following this filtering approach, we obtained 8764 SNPs in 517 individuals for further analyses. Because the adults had on average more missing data than the hatchlings, we generated a second dataset comprising only the 82 wild-collected adults, filtering with the same criteria as above. This second dataset contained 4943 SNPs and was used to infer the number of genetic lineages present. All downstream analyses were conducted in R 4.0.3 (R Core Team, 2021). We read the vcf file using the R package vcfR v1.8.0 (Knaus & Grünwald, 2017).

We amplified and sequenced an 814 bp fragment of the COI mitochondrial gene for all adults in our dataset, plus reference *T. douglasi* and *T. poppense* from other populations (Table S2) using a nested PCR. We first amplified DNA with primers 3’-TCCAATGCACTAATCTGCCATATTA-5’ and 3’-GGAACNGGATGAACAGTTTACCCNCC-5’. We then amplified a subset of this fragment using the former and 3’-CAACATTTATTTTGATTTTTGG-5’. All primer sequences were obtained from Simon et al., (1994). All reactions were conducted in 20 μL of Promega Buffer 1X containing 0.3 μM of each primer, 0.2 mM of each dNTP and 0.02 U/μL G2 Taq polymerase (Promega). All cycles consisted of 5mn at 94°C followed by 20 (first PCR) and 35 (nested PCR) cycles of 30 s at 95°C, 1 mn at 50°C, 1 mn at 72°C, and a final extension at 72°C for 10 mn. The PCR products were sequenced both ways at Microsynth (Switzerland), and we generated the consensus from the forward and reverse sequences using BioEdit.

### Population structure and phylogeny

To identify the number of genetic lineages present across the two transects, we conducted a clustering analysis on the adults using faststructure v1.0 (Raj et al., 2014), with K ranging from 1 to 10. We then used the chooseK.py script to select the value of K that maximises marginal likelihood.

We also wanted to assess whether individuals collected along the two transects belonged to known lineages of the parthenogenetic species *T. douglasi* or the sexual species *T. poppense*. Although we did not expect populations of the sexual species *T. poppense* in the surveyed geographic range (Law & Crespi 2002), we could not exclude the presence of geographically isolated *T. poppense* populations. This is because *T. douglasi* is very closely related *T. poppense* and females of these two species cannot be distinguished morphologically (Vickery & Sandoval, 1999). We therefore used reference individuals for species assignments. These reference individuals, five *T. douglasi* and 26 *T. poppense* individuals from different populations (Table S2) were identified in previous studies (Jaron et al., 2022; Schwander et al., 2011), with DNA extractions still available in the lab. We used these 31 reference individuals to study their phylogenetic relationship with the genetic lineages identified in our newly sampled populations for mitochondrial and nuclear (RADseq) markers. We generated RADseq and COI data for these individuals with the same protocols as described above. For the RADseq data, we retained genotype calls with a minimum depth of 8 instead of 11, as allelic dropout would cause less bias for phylogenetic reconstruction than for inference of reproductive modes (largely based on heterozygosity estimations, see below). We discarded loci that had a minor allele count lower than three and were genotyped in less than 80% of all individuals. This dataset contained 16391 sites. The COI dataset contained 159 polymorphic sites. Note that some individuals had multiple COI copies (as indicated by double peaks in the sequencing trace), as expected for the presence of nuclear copies of mitochondrial genes or heteroplasmy. In these cases, we called the allele corresponding to the largest peak, and used an ambiguous genotype following the IUPAC code when both peaks were equally large. We added the COI sequence from one individual of *T. cristinae* as an outgroup (EU251511.1; (Schwander et al., 2011). We then aligned the sequences using the MUSCLE algorithm (Edgar, 2004). We reconstructed separate maximum likelihood phylogenies for the nuclear and mitochondrial data, with iqtree2 (Minh et al., 2020), using ModelFinder Plus to select the best substitution model in each case. We used the cophylo function of the R package phytools (Revell, 2012) to plot them as a cophylogeny.

### Reproductive mode analyses

For individuals of the Manchester and Orr transects, we calculated heterozygosity as the proportion of the SNPs that were heterozygous. Note that because we only retained polymorphic positions in our dataset, this measure overestimates genomic heterozygosity and is hereafter referred to as “relative heterozygosity”. We observed that relative heterozygosity increased linearly with the proportion of missing data (Figure S1). This increased heterozygosity is likely due to merged paralogs, which are enriched in individuals with more missing data as paralogues have higher coverage than single-copy loci and are therefore preferentially retained (Supplementary Material). We therefore corrected relative heterozygosity to remove the effect of missing data by using the residuals of the linear regression and adding the intercept value (Figure S1).

To assess whether hatchlings from eggs laid after mating were produced via sex or parthenogenesis, we used two complementary measures. For the first measure, we looked for genotypes in hatchlings that cannot be produced via parthenogenesis. At loci where the mother is homozygous, offspring can only be heterozygous if they were produced from a fertilized egg (in the absence of genotyping errors; see also Brown et al., 2021). For each offspring, we therefore computed the proportion of heterozygous loci that had a homozygous genotype in the mother. Loci at which either the mother or the offspring were not genotyped were excluded.

For the second measure to assess whether hatchlings were produced via sex or parthenogenesis, we compared relative heterozygosity between mothers and their offspring. We expected heterozygosity levels of hatchlings produced via sex to be similar to that of the mother, or higher in hatchlings sired by a male originating from a different lineage than the mother. On the other hand, we expected heterozygosity levels of parthenogenetically produced hatchlings to be lower than that of their mother. A decrease is expected because facultative parthenogenesis generally occurs via automixis (Suomalainen et al., 1987). Under automixis, meiotic divisions occur and diploidy is restored secondarily via fusion or duplication of meiotic products. Similar to selfing, automixis can result in a loss of heterozygosity between mothers and offspring (Pearcy et al., 2006; Suomalainen et al., 1987). However, heterozygosity losses can only be observed at sites where the mother was heterozygous, and where heterozygosity losses are not mechanistically constrained or removed by selection (Jaron et al., 2022). The extent of the heterozygosity loss then depends on the type of automixis and the amount of recombination (Pearcy et al 2006, Suomalainen et al 1987). In the most extreme case (gamete duplication) a haploid gamete duplicates to produce a fully homozygous zygote. Other forms of automixis (e.g., terminal or central fusion) result in a partial loss of heterozygosity. In these cases, offspring heterozygosity depends on the heterozygosity of the mother, and on the position and frequency of recombination events. By combining the two heterozygosity-based measures, we were able to unambiguously assign each of the 365 genotyped hatchlings from eggs produced after mating to either sexual or parthenogenetic offspring (see Results).

We then asked whether females from different genetic lineages used different reproductive strategies. To answer this question, we tested whether there were significant differences between lineages in (i) hatching success of eggs laid before mating, (ii) hatching success of eggs laid after mating, and (iii) the proportion of post-mating eggs that were fertilized (which measures the propensity to reproduce via sex) using binomial Generalized Linear Models (GLMs), tested their significance using the Anova function of the R package car v3.0-10 (Fox & Weisberg, 2019) and performed multiple comparisons using the glht function implemented in the R package multcomp v1.4-16 (Hothorn et al., 2008). We also looked for differences in hatching success of eggs before vs after mating within each lineage using binomial GLMMs implemented in the R package lme4 (Bates et al., 2015), using the mother identity as random factor.

In order to test whether variable population sex ratios along the Orr and Manchester transects were generated by sex ratio distortion, we quantified the sex ratio in sexually produced offspring. Because female and male hatchlings cannot be distinguished morphologically, we sexed offspring genetically by combining heterozygosity at X-linked loci and the read depth ratio between the X-linked and autosomal loci. Because *Timema* males have only one copy of the X chromosome and no Y (XX:XO sex determination; Schwander & Crespi, 2009), there should be no heterozygosity for X-linked loci. Furthermore, X-linked loci should have half the coverage compared to autosomal loci in males, but similar coverage in females. To sex sexually produced offspring via these approaches, we first had to identify X-linked scaffolds in the *T. douglasi* reference genome (Jaron et al., 2022).

To identify X-linked scaffolds we used a combination of available genomic data from *T. douglasi* females (Bioproject accession number: PRJNA670663) in addition to whole-genome sequencing of two male *T. douglasi*. DNA extractions were done on adult male carcasses using the Qiagen Mag Attract HMW DNA kit following the manufacturer instructions. Sequencing libraries were generated for each male using a TruSeq DNA nano prep kit (550bp insert size). Libraries were then sequenced (150b, paired-end) using an Illumina HiSeq 4000 at the Lausanne Genomic Technologies Facility. Reads were trimmed before mapping using Trimmomatic v0.36 (Bolger et al., 2014) to remove adapter and low-quality sequences (options: ILLUMINACLIP:3:25:6 LEADING:9 TRAILING:9 SLIDINGWINDOW:4:15 MINLEN:90). Reads from each individual were then mapped to the reference genome using BWA-MEM v0.7.15 (Li & Durbin, 2010). Multi-mapping and poor quality alignments were filtered (removing reads with XA:Z or SA:Z tags or a mapq < 30). PCR duplicates were removed with Picard (v. 2.9.0) (http://broadinstitute.github.io/picard/). Coverage was then estimated for all scaffolds at least 1000 bp in length using BEDTools v2.26.0 (Quinlan & Hall, 2010). To compare coverage between males and females, coverage was first summed up for all male and all female libraries per scaffold. Male and female coverage was then normalised by modal coverage to adjust for differences in overall coverage. X-linked scaffolds were then identified using the log2 ratio of male to female coverage. Scaffolds were classified as X-linked if the log_2_ ratio of male to female coverage was within 0.1 of the value of the X-linked peak (Figure S3), and as autosomal otherwise.

By distinguishing between autosomal and X-linked loci, we then sexed sexually produced offspring from our controlled crosses using the RADseq data. We first standardised read depth per individual, dividing the depth value at each locus by the average depth for that individual. We then computed the X to autosomes depth ratio by dividing average standardised depth of loci on the X chromosome by that of autosomal loci (see supplementary material). Using this depth ratio in combination with heterozygosity on the the X chromosome, we were able to sex 176 out of the 211 sexually produced offspring (see Figure S5; the remaining 35 offspring had insufficient sequencing coverage for these analyses). We tested for deviations from a 50:50 sex ratio among sexually produced offspring overall using a chi-squared test. We further investigated whether there were deviations within each family. For this, we looked whether the 95% binomial proportion confidence interval of the observed sex ratio overlapped with the expected 50:50 sex ratio. In addition, we conducted Fisher’s exact tests for each family.

## Results

We found that adults from geographically close populations differed strongly in their sex ratio, with significant sex ratio variation among populations overall (Figures 1A, 1C; Manchester: Likelihood ratio χ2 = 456.49, df = 11, p < 2 x 10-16; Orr: Likelihood ratio χ2 = 106.46, df = 11, p < 2 x 10-16). The Orr transect consisted of a mosaic of female-biased and female-only populations. Five populations (Orr 1, 5, 8, 9 and 17) had between 77.8% and 91.7% of females, while the others had only females (Figure 1A). By contrast, we found an abrupt transition from roughly even to strongly female-biased or female-only sex ratios along the Manchester transect. Sex ratios in populations 1 to 7 ranged from 43.2% to 54.7% of females, while populations 8 to 12 consisted of at least 93.5% females (Figure 1C).

We then investigated whether the field-collected individuals belonged to different genetic lineages using a clustering analysis. The best supported number of lineages among all genotyped field-collected individuals was three. We found a gradient of genotype incidences along the transects. The first lineage (blue in Figure 1B) was mostly found along the Orr transect, with only one individual in Manchester 10, and is hereafter referred to as the “Orr lineage”. The second lineage (yellow in Figure 1B) was found mostly in the eastern section of Manchester and is hereafter referred to as the “eastern Manchester lineage”. Finally, the third lineage (red in Figure 1B) was mainly found in the western section of Manchester and is hereafter referred to as the “western Manchester lineage”. We found little evidence for admixture between lineages in 14 out of the 15 populations tested, with only one female from Orr 11 (17-2162) appearing to be a hybrid, perhaps triploid, between the three lineages (Figure S2). In one population however, Manchester 2, all 5 genotyped individuals appeared to be admixed, with approximately equal contributions of the eastern and western Manchester lineages. These results were robust with varying numbers of clusters (k in faststructure), indicating that these were most likely hybrids and not a fourth lineage (Figure S2).

In our nuclear phylogeny, all individuals collected along the two transects formed a monophyletic clade with the parthenogenetic T. douglasi reference individuals, indicating that all three lineages belong to T. douglasi. All T. poppense reference individuals formed a second, monophyletic clade (Figure 1E). The mitochondrial phylogeny also showed two monophyletic groups. One contained mostly T. poppense, with five T. douglasi individuals (including four from our transects). The second contained mostly T. douglasi, with three T. poppense individuals (Figure 1F). Individuals belonging to the eastern Manchester lineage formed two distinct mitochondrial subclades, which were not segregated geographically.

We isolated 352 females to characterise their reproductive mode and assess the hatching success of the eggs they laid as virgins. 265 of these females laid at least five eggs as virgins. At least one unfertilized egg hatched from the clutches produced by 234 (88.3%) of them, indicating that they were able to reproduce via parthenogenesis. In addition, females capable of parthenogenesis were found in all populations tested. The hatching success of unfertilized eggs varied between 3.3 and 100% (Figure 2A).

**Figure 2.**
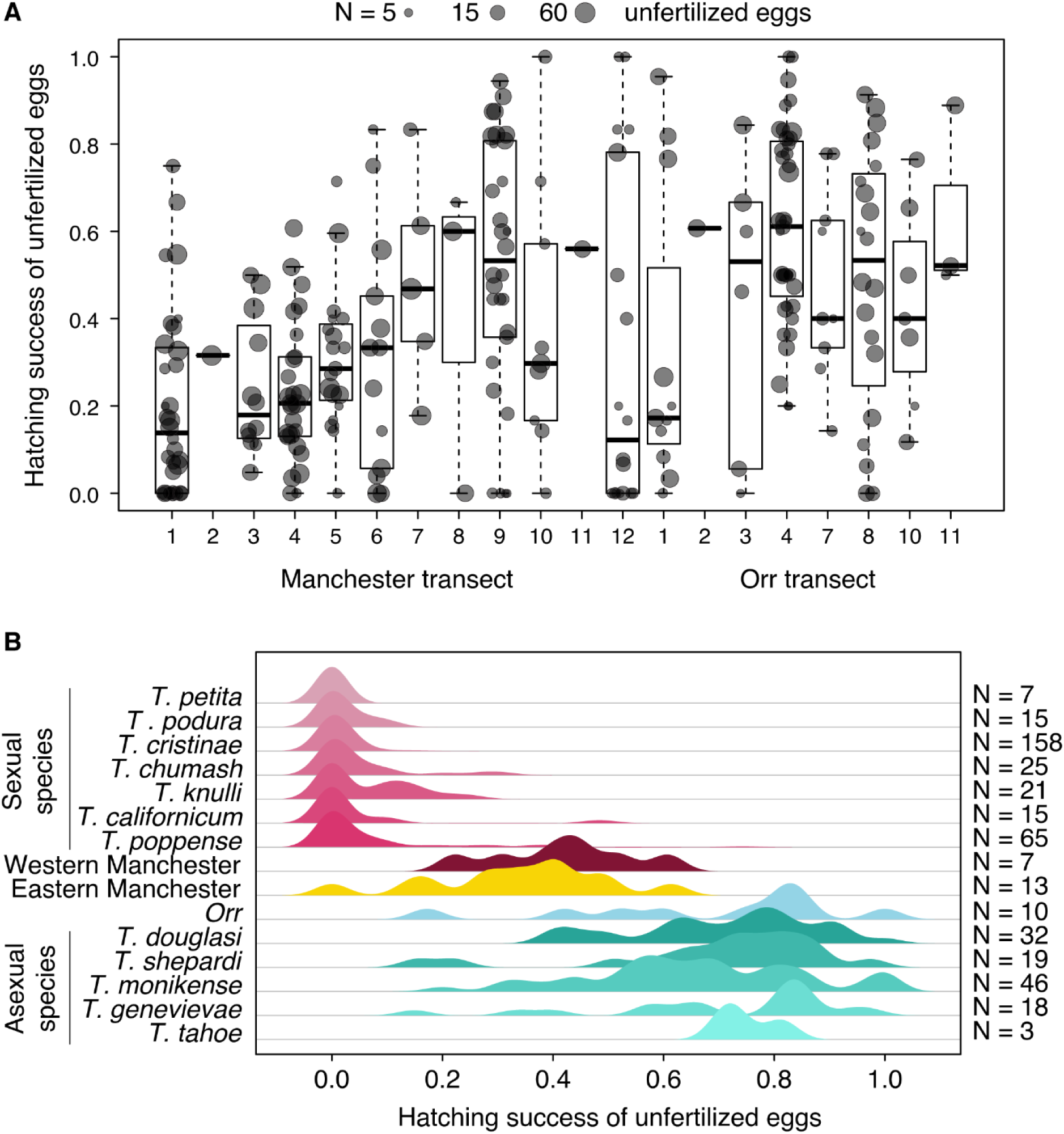
Parthenogenesis capacity (**A**) in field-collected females from different populations along the Manchester and Orr transects and (**B**) in females from different obligate sexual and obligate asexual species, measured by the hatching success of unfertilized eggs. Dot size in **A** is proportional to the number of unfertilized eggs laid by each female. In **B**, N refers to the number of females for which hatching success was measured.

The hatching success of unfertilized eggs from the Orr lineage was within the range of obligate asexual *Timema* species, while for the two Manchester lineages, it was intermediate between obligate sexuals and asexuals (Figure 2B).

In order to gain insights into the mechanisms of automictic parthenogenesis in females from the Orr and Manchester transects, we compared relative heterozygosity in mothers and their offspring produced from eggs laid prior to mating. Heterozygosity losses can only be observed at sites where the mother was heterozygous, and where heterozygosity losses are not mechanistically constrained or removed by selection (Jaron et al., 2022). However, if heterozygosity loss does occur, the extent of it can inform on the type of automixis (Pearcy et al 2006). We found extensive loss of heterozygosity or extremely low relative heterozygosity in all offspring from eggs laid prior to mating (Figure 3), as would be expected from gamete duplication or terminal fusion without recombination. With one exception, relative heterozygosity in these offspring amounted to 2 - 3% and was not correlated with their mothers’ heterozygosity (as would be expected i.e., under central or terminal fusion automixis with recombination). In addition, most, if not all of the heterozygosity observed in hatchlings is caused by false positives (see Supplementary Material), meaning that these hatchlings are most likely completely homozygous. Only one parthenogenetic hatchling from female 17-1793 retained some heterozygosity (Figure 3), which could suggest that it was produced by a different parthenogenesis mechanism than the other individuals.

**Figure 3.**
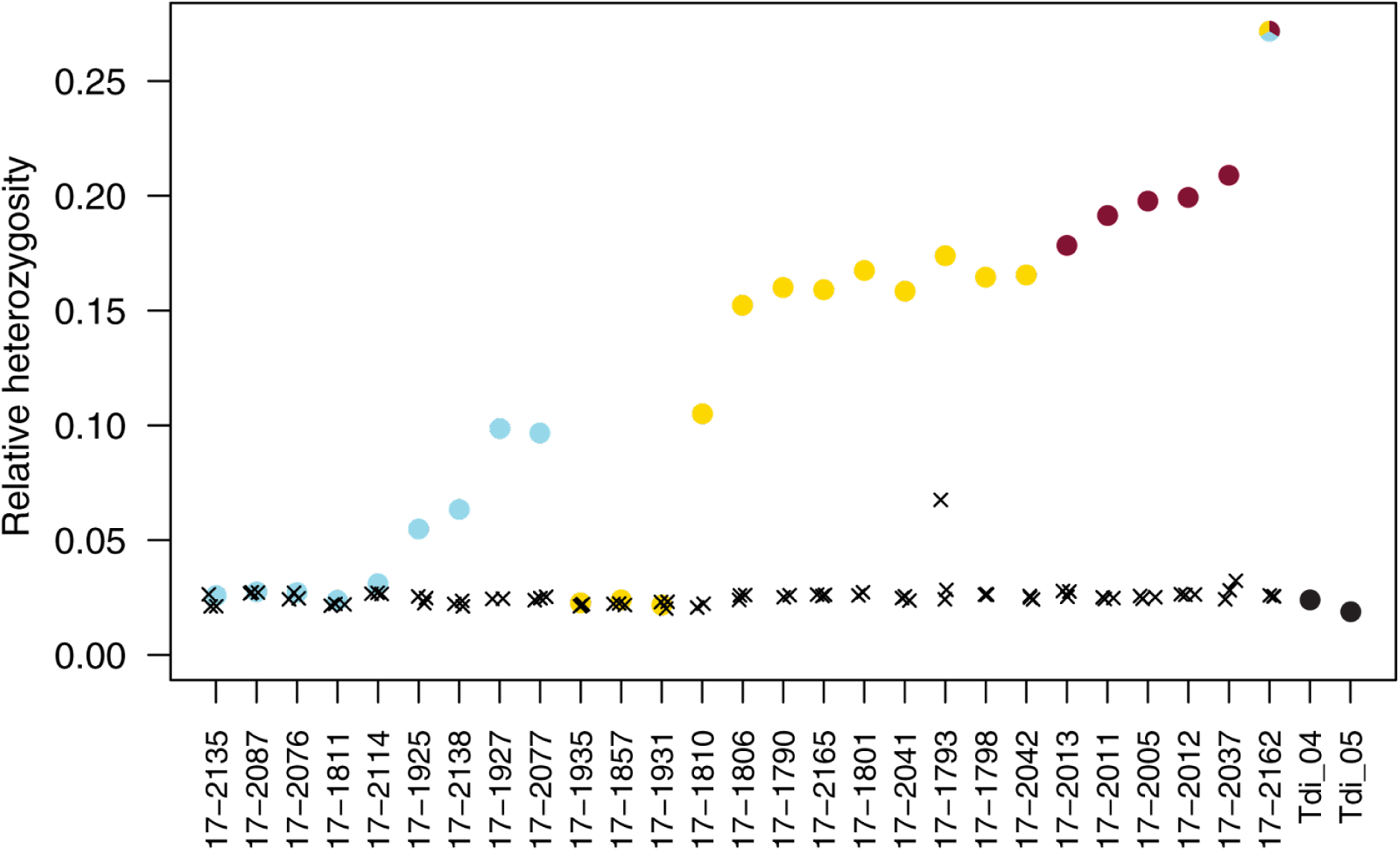
Relative heterozygosity of mothers (circles coloured as in Figure 1) and their offspring produced from unfertilized eggs laid before mating (black crosses, two to three per mother), and of two reference parthenogenetic individuals (Tdi_04 and Tdi_05) from Jaron et al. (2022).Note the much higher heterozygosity in one hatchling from family 17-1793, indicating that it could have been produced via a different mechanism of parthenogenesis.

We were able to unambiguously infer whether each offspring from eggs laid after mating was produced via sex or parthenogenesis by using a combination of relative heterozygosity and proportion of genotype transitions from homozygous in the mother to heterozygous in the offspring (Figure 4; see also Supplementary Material). These inferences revealed that 211 (58.3%) of offspring were produced by sex, and 151 (41.7%) were produced via parthenogenesis.

**Figure 4.**
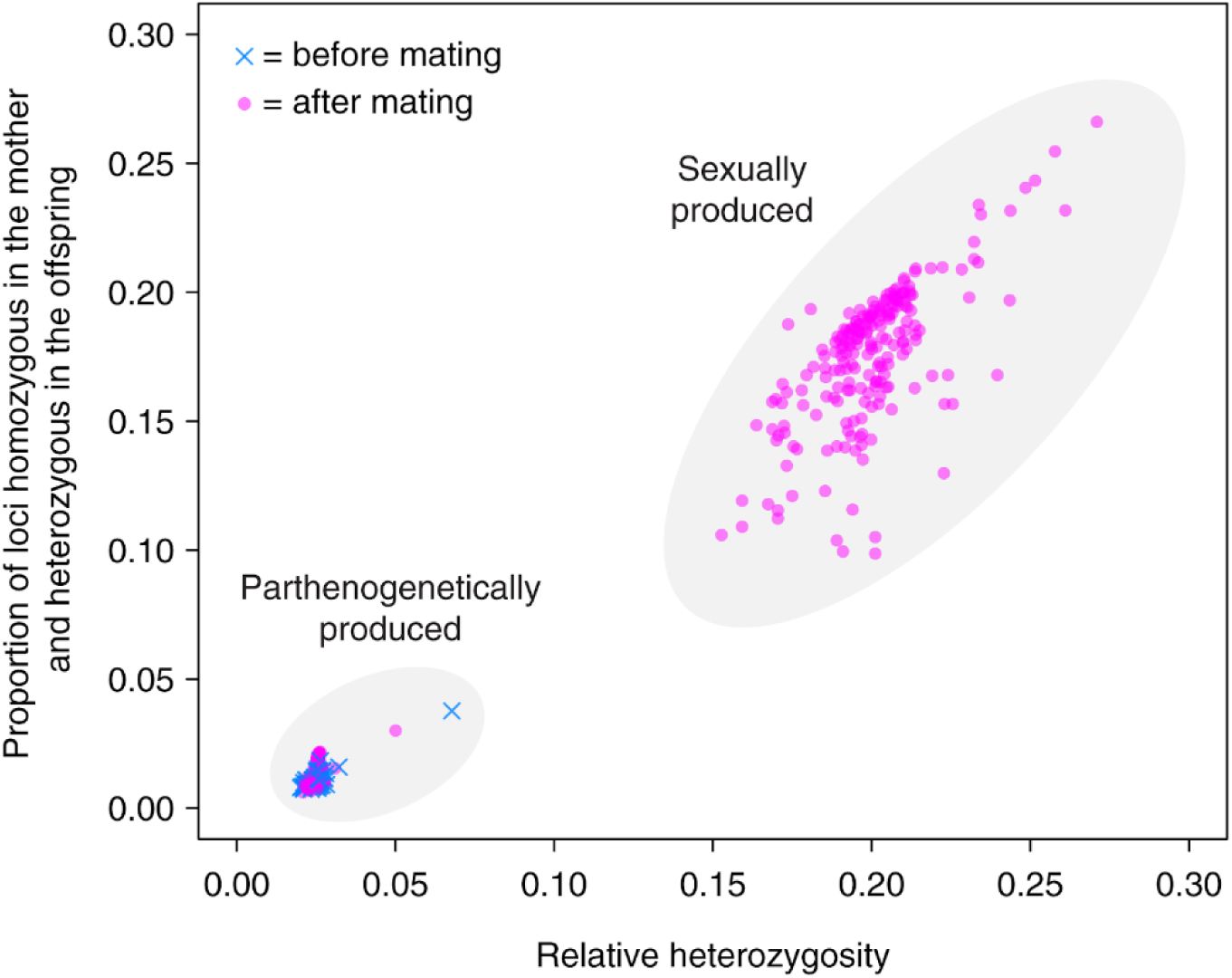
Relative heterozygosity and proportion of transitions from homozygous genotype in the mother to heterozygous in the offspring, in offspring from eggs laid before (blue crosses) and after mating (pink dots). These data were used to infer whether offspring from eggs laid after mating were produced via sex or parthenogenesis.

We then looked at differences in the parthenogenetic capacity or the propensity to use sex following mating between females from the Orr, western and eastern Manchester lineages. All 32 females for which we were able to infer the lineage via genotyping were capable of parthenogenesis, but with varying efficiency (Figure 5A). Note that one of these females (17-1937 from the eastern Manchester lineage) did not produce any offspring from the 13 eggs she laid before mating, but she was capable of parthenogenesis since all three offspring produced from the 14 eggs laid after mating were produced via parthenogenesis. Hatching success of eggs laid before mating was on average higher for females of the Orr lineage (67%) than of the two other lineages (western Manchester: 42%, *z* = 2.51, *p* = 0.032; eastern Manchester: 33%, *z* = 3.53, *p* = 0.001; Figure 5A). There was no significant difference between the lineages in hatching success of eggs laid after mating (χ^2^ = 0.003, *df* = 2, *p* = 0.99; Figure 5B). Mating influenced the egg hatching success in all three lineages. In the eastern and western Manchester lineages, eggs laid after mating had a significantly higher hatching success than those laid by virgin females (western Manchester: 55% vs 42%, *z* = 2.0, *p* = 0.04; eastern Manchester: 55% vs 33%, *z* = 4.7, *p* = 2.7*10^-6^). On the other hand, in the Orr lineage, eggs laid after mating hatched at a lower rate than those laid before mating (56% vs 69%, *z* = -4.7, *p* = 2.7*10^-6^). We also found that the proportion of eggs laid after mating that were actually fertilized was highly variable among females. Twelve females fertilized between 80% and 100% of their eggs laid after mating, while eleven fertilized none of them. Only five fertilized an intermediate proportion (14% to 50%). There were significant differences in the proportion of fertilized eggs between lineages. Females from the Orr lineage fertilized on average fewer eggs (10%) than those from the eastern and western Manchester lineages (79%, *t* = 4.0, *p* = 0.0002 and 95%, *t* = 3.5, *p* = 0.001 respectively). All females from the western Manchester lineage fertilized more than 80% of their post-mating eggs, while seven out of 10 females from the Orr lineage fertilized none of them. Finally, the eastern Manchester lineage consisted mostly of females with either a very high or very low fertilisation frequency (Figure 5C). The variation in fertilization frequency was not explained by the two mitochondrial subclades. The putatively triploid hybrid female (17-2162) was also capable of facultative parthenogenesis: a large portion of eggs laid prior to mating hatched (24 out of 27), and out of the 17 genotyped hatchlings from the eggs she laid after mating, seven were produced via sex and 10 via parthenogenesis.

**Figure 5:**
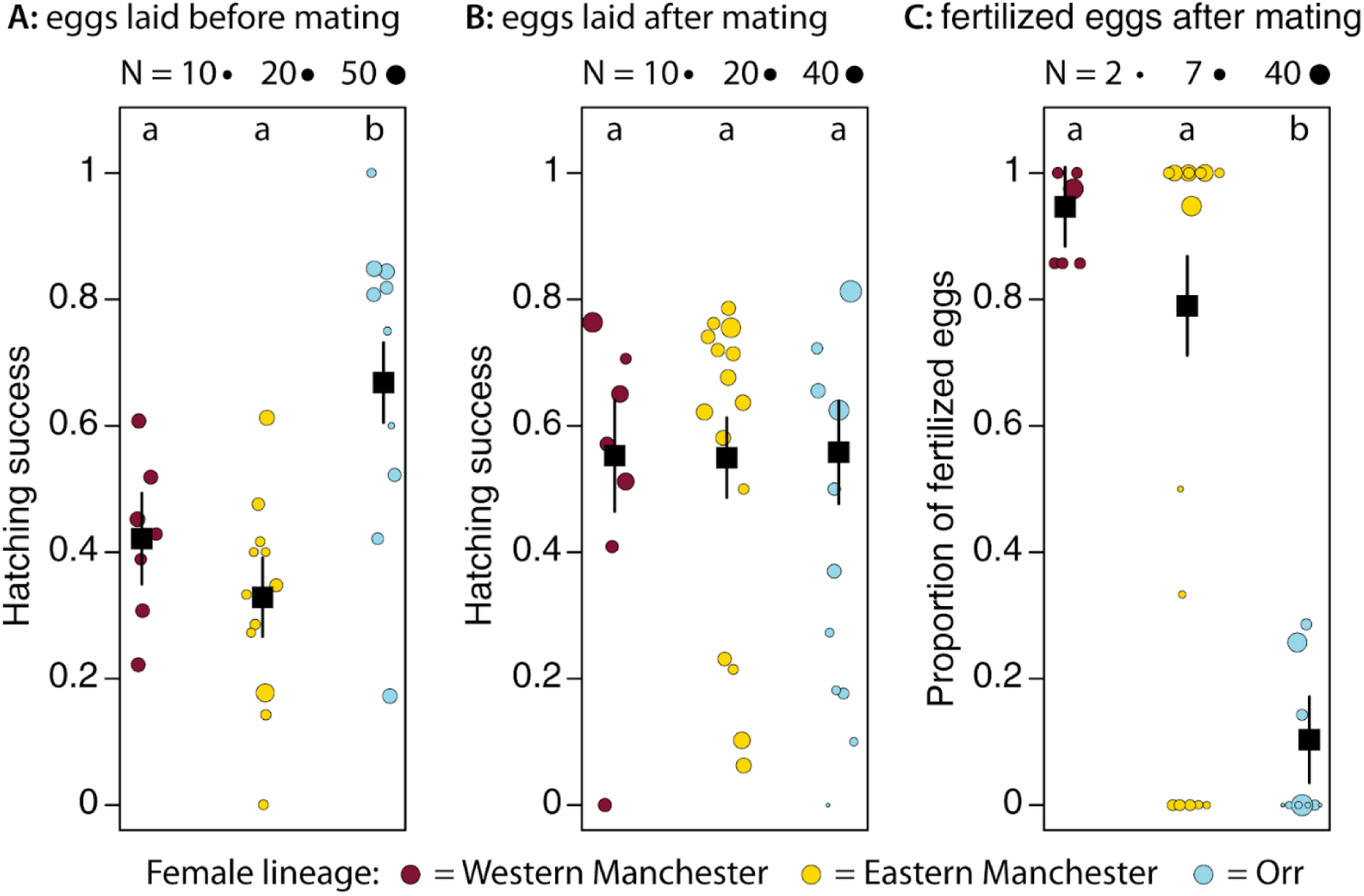
Reproductive mode differences between females of the three genetic lineages (distinguished by colours as in Figure 1). **A**: Hatching success of eggs laid prior to mating, which measures the capacity to do parthenogenesis. Dot size proportional to the number of eggs laid before mating. **B**: Hatching success of eggs laid after mating. Dot size proportional to the number of eggs laid after mating. **C**: The proportion of sexually produced hatchlings that originated from eggs laid after mating, which measures the propensity to reproduce via sex. Dot size proportional to the number of hatchlings genotyped to assess whether they were produced via sex or parthenogenesis. In all panels: the black squares and bars represent the weighted average per lineage +/- standard error. Different letters indicate significantly different groups (*p* < 0.05 in multiple comparison tests after binomial GLMs).

Finally, we tested whether sex ratio distortion could be contributing to female-biased sex ratios by looking at the sex ratio of sexually produced offspring. A driving X chromosome should lead to an excess of female offspring. Across all families, there were 81 female and 95 male offspring, providing no evidence for biased sex ratios (χ^2^ = 1.1, *p* = 0.29). The same was observed within families (all *p* > 0.5; Figure S6).

## Discussion

How transitions between fundamentally different reproductive modes – such as sex and obligate parthenogenesis – can occur, is unclear. Transitions might be abrupt, with a direct switch from sex to obligate parthenogenesis within one generation. Alternatively, transitions might be gradual and involve a period where individuals are capable of facultative parthenogenesis (Schwander & Crespi, 2009 Neiman et al., 2014). In this study, we discovered high frequencies of males in populations of *Timema douglasi*, a species previously believed to reproduce solely via obligate parthenogenesis. These populations were characterized by variable sex ratios, which were caused by different frequencies of sexual versus parthenogenetic reproduction among facultative parthenogens, and not by sex ratio distortion. These findings raise the possibility that obligate parthenogenesis, which evolved repeatedly in the genus *Timema*, may generally derive from facultative parthenogenesis, with facultative parthenogenesis being a transient state. Two genetically differentiated lineages of *T. douglasi* reproduced through true facultative parthenogenesis, with good hatching success of unfertilized eggs but a high proportion of sexually produced offspring when mated. They were found geographically close to a third lineage of the same species (Orr), in which females had a high capacity for parthenogenesis and mostly did not fertilize their eggs when mated. Furthermore, eggs produced after mating by females of this lineage suffered from a lower hatching success, suggesting that fertilisation of eggs that would normally develop into parthenogenetic offspring can sometimes result in developmental failure. These results may point to a tradeoff between sexual and parthenogenetic pathways as was previously suggested to be the case in sexual *Timema* populations: the ability for spontaneous parthenogenesis likely reduces hatching success of fertilized eggs (Schwander et al., 2010). However, lower hatching success of post-mating eggs could also be due to females being older. This effect would be compensated by the positive effect of fertilization on egg hatching success in both Manchester lineages.

Females of the two facultatively parthenogenetic lineages from the Manchester transect had a lower capacity for parthenogenesis than those of the largely obligate one and featured different propensities to reproduce sexually following mating. Females of the first lineage (western Manchester) generally fertilised all their eggs when mated, which resulted in higher hatching success than under parthenogenesis. Females of the second lineage (eastern Manchester) were more variable. Approximately half (8 out of 15) of them reproduced the same way as the western Manchester lineage: using mainly sex when mated but being able to reproduce via parthenogenesis. Five females reproduced only via parthenogenesis, similarly to females of the Orr lineage. The two remaining females seemed to use both sex and parthenogenesis at approximately equal frequencies (facultative parthenogenesis), but genotyping larger clutches would be needed to assess whether this was indeed the case.

The mechanisms that maintain reproductive polymorphism (facultative and largely obligate parthenogenesis) within populations, and those that cause the sharp transition from equal to strongly female-biased sex ratios over a few hundred meters along the Manchester transect remain an open question. Since more than half of the females from the eastern Manchester lineage were able to reproduce via sex and fertilized most or all of their eggs following mating, a male immigrating from a nearby sexual population would be expected to have extremely high reproductive success. This should then lead to an increase in male frequency in the next generation. One possible explanation for the maintenance of different sex ratios between neighbouring populations is that dispersal is very limited in these wingless insects (Sandoval, 2000), and that it is just a matter of time until the female-biased Manchester populations shift towards more equal sex ratios. However, variable selection pressures along the transect could also play a role, favouring respectively sexual or parthenogenetic reproduction (Burke & Bonduriansky, 2018). This could help explain why males, even though present in small numbers in the strongly female-biased populations (such as Manchester 12 with <1% males), have not triggered sex ratio shifts. Identifying such selection pressures would constitute a major leap towards understanding the costs and benefits of different reproductive modes in natural populations.

Our comparison of heterozygous genotypes in mothers and their parthenogenetically produced offspring also provides insights into the mechanisms of parthenogenesis and the evolution of obligate parthenogenesis in *Timema*. Relative heterozygosity in all but two parthenogenetic offspring (out of 228 produced before or after mating) was around 2.5%, regardless of their mothers’ heterozygosity. Given the 2.5% heterozygosity is most likely generated by structural variation between individuals and divergence between paralogs (Supplementary Material), the main mechanism of parthenogenesis in *Timema* thus appears to generate genome-wide homozygosity in a single generation. This would be expected under specific forms of automixis, notably under gamete duplication or fusion of non-recombined sister chromatids (i.e., terminal fusion without recombination). The two remaining out of 228 parthenogenetic offspring (one originating from an unfertilized egg laid prior to mating, the other from an unfertilized egg laid after mating; see Figure 4), produced by different females, had somewhat higher heterozygosity. This suggests that some females are able to reproduce via two different parthenogenesis mechanisms in addition to sex, as was described for spontaneous parthenogenesis in sexual *Daphnia* (Svendsen et al., 2015). Automixis with “complete” heterozygosity loss in a single generation is also the proposed mechanism of parthenogenesis in obligate asexual *Timema* species (Jaron et al., 2022) and of spontaneous parthenogenesis in sexual *Timema* species (Schwander & Crespi, 2009). In *Timema*, obligate parthenogenesis thus likely evolved via a gradual increase of the capacity for parthenogenesis, with a conserved proximate mechanism.

Our finding of facultative parthenogenetic lineages of *T. douglasi* questions the directionality of transitions between reproductive modes in this genus and beyond. So far, sex with low levels of tychoparthenogenesis was believed to be the ancestral state in *Timema*, followed by repeated transitions to obligate parthenogenesis (Schwander & Crespi, 2009). However, our results open the possibility that facultative parthenogenesis be an ancestral state in this genus, and that the different lineages have transitioned towards more obligate strategies. This would require that obligate parthenogenesis has evolved several times in the *T. poppense/douglasi* species complex. Phylogenetic analyses of the *Timema* genus revealed that *T. douglasi* consisted of multiple lineages that independently derived from the sexual species *T. poppense* (Schwander et al., 2011). Repeated transitions towards parthenogenesis are not surprising if the ancestor of these species was already capable of reproducing via facultative or spontaneous parthenogenesis, and if selection has the opportunity to increase parthenogenetic capacity, for example due to mate limitation (Schwander et al., 2010).

Alternatively, the facultative parthenogenetic *T. douglasi* lineages could represent a re-evolution of sex. Sex could in theory reappear in all-female lineages via accidental male production. Rare male production in species with XX/X0 sex determination such as *Timema* can occur via aneuploidy at the X chromosome, because the resulting X0 individuals would develop into males (Pijnacker & Ferwerda, 1980). However, such a re-expression of the sexual pathway would require the molecular bases underlying the sexual functions of both sexes to be maintained. Re-expression of sex is therefore possible at least shortly after the transition to parthenogenesis, which may also have been the case in the stick insects *Clitarchus hookeri* (Morgan-Richards et al., 2019) and *Bacillus rossius* (de Vichet, 1944). Occasional male production and re-expression of the sexual pathway in otherwise obligate parthenogens paves the way for cryptic sex, which could contribute to explaining their long-term persistence (Freitas et al., 2023).

Finally, gene flow between sexual and parthenogenetic lineages could contribute to frequent transitions between reproductive modes. One such example is contagious parthenogenesis, whereby rare males from otherwise parthenogenetic lineages transmit the capacity for parthenogenesis to a related sexual lineage via hybridization. This has been documented in some aphids (Delmotte et al., 2001), wasps (Sandrock & Vorburger, 2011) and *Daphnia* (Innes & Hebert, 1988). Conversely, gene flow from a sexual to mostly but not completely parthenogenetic species could in theory result in an increase of the frequency of sex in the latter. Occasional gene flow between *T. poppense* and *T. douglasi* is suggested by the mismatches we observed between nuclear and mitochondrial genomes in about 10% of our individuals. This could be caused by rare hybridization events between the two species in both directions and contribute to the maintenance of different reproductive modes within populations. Formal tests of gene flow between *T. poppense* and *T. douglasi* would shed light onto this question.

More generally, the faculty for occasional and facultative sex in *T. douglasi*, a species previously believed to be obligately parthenogenetic, adds to recent re-evaluation of the obligate status of some parthenogenetic species (Boyer et al., 2021; Kuhn et al., 2021; Laine et al., 2022). Many such “ancient asexuals” could in fact have retained the capacity for occasional, cryptic sex (Schurko et al., 2009). They would thus represent extremes of a continuum from mostly sexual to mostly parthenogenetic reproduction. On the other end of this continuum, many taxa that were thought to reproduce solely via sex were recently found to be capable of parthenogenesis, though often at anecdotal frequency and with limited success (e.g. Booth et al., 2012; Fields et al., 2015; Groot et al., 2003; Ryder et al., 2022, but see Kratochvíl et al., 2020).

Whether a transient facultative state is a frequent feature in transitions between extremes of the reproductive continuum still remains to be investigated. Still, the factors that would favour obligate over more facultative strategies remain elusive. Facultative parthenogenesis is believed to be an efficient reproductive mode, combining the “best of both worlds” – the long-term advantages of recombination and segregation along with the short-term advantages of asexuality. However, the loss of sex in facultative parthenogens could in theory be driven by sexual conflict (if mating always reduces female fitness; (Burke & Bonduriansky, 2017). Alternatively, the lower parthenogenesis capacity observed in the facultatively asexual lineages than in both the more obligate lineage (Orr) and the obligate asexual species, as well as interspecific comparisons in facultatively parthenogenetic mayflies (Liegeois et al., 2021) suggest that the efficiency of sex and parthenogenesis are traded-off against each other, consistent with the “jack of all trades, master of none” hypothesis. In this case, whether obligate strategies are likely to replace facultative ones will depend on local ecological conditions favouring sex or parthenogenesis, and on the fluctuations of such conditions.

In conclusion, we discovered the first case of facultative parthenogenesis in *Timema* stick insects, a genus where obligate parthenogenesis has evolved repeatedly from a sexual ancestor, with an apparently conserved proximate mechanism of parthenogenesis. We found three genetic lineages that differed both in their capacity to reproduce via parthenogenesis and in their propensity to reproduce sexually. It is thus far unclear whether these populations reverted to sex starting from a largely asexual ancestor, or they are sexual relicts in the largely asexual *T. douglasi*, where all other populations have transitioned to obligate parthenogenesis already. The latter would suggest that transitions to asexuality were numerous during the evolution of this species, and that asexual populations almost always prevailed on sexuals. In that case, we could be witnessing an ongoing transition to asexuality via facultative parthenogenesis. Why facultative parthenogenesis would be a transient stage in *Timema* and which factors would favour a transition to obligate parthenogenesis would thus be exciting foci for future research.

## Acknowledgements

We thank Ian S. Ford and Armand Yazdani for their great help in the field, Karine Légeret for her assistance in the daily monitoring of egg laying and hatching in the lab, and Zoé Dumas and Marjorie Labédan for wet lab help. Comments by three anonymous reviewers improved the manuscript. Most analyses were run using the computing infrastructure at the University of Lausanne maintained by the former Vital-IT platform of the SIB Swiss Institute of Bioinformatics (SIB) and the DCSR. Preprint version 4 of this article has been peer-reviewed and recommended by Peer Community In Evolution (https://doi.org/10.24072/pci.evolbiol.100551).

## Data, scripts, code, and supplementary information availability

All demultiplexed RAD-seq reads have been deposited in NCBI’s sequence read archive under BioProject ID PRJNA804475 and whole genome sequence reads of T. douglasi males under BioProject ID PRJNA808673. COI sequences have been deposited on GenBank under accessions OP379833 – OP379903. The code used to run the analyses presented in this article as well as the supplementary text and figures are available on GitHub under https://github.com/glavanc1/Timema_facultative_partheno, archived in Zenodo under https://zenodo.org/badge/latestdoi/266724954.

